# Apparent timescaling of fossil diversification rates is caused by sampling bias

**DOI:** 10.1101/2024.09.11.612481

**Authors:** Bouwe R. Reijenga, Roger A. Close

## Abstract

Negative scaling relationships between both speciation and extinction rates on the one hand, and the age or duration of organismal groups on the other, are pervasive, and recovered in both molecular phylogenetic and fossil time series. The agreement between molecular and fossil data hints at a universal cause, and potentially to incongruence between micro- and macroevolution. However, the existence of negative rate scaling in fossil time series has not undergone the same level of scrutiny as in molecular data. Here, we analyse the marine fossil record across the last ~538.8 Ma of the Phanerozoic to investigate the presence and strength of negative rate scaling. We find that negative rate scaling arises under commonly applied age range-based per-capita rates, which do not control for sampling bias, but are severely reduced or absent when metrics are used that do correct for sampling. We further show by simulation that even moderately-incomplete sampling of species occurrences through time may induce rate scaling. We thus conclude that there are no significant scaling relationships present in these fossil clades, and that any apparent trend is caused by sampling artefacts and taxonomic practices. If rate scaling in molecular phylogenies is genuine, the absence of such a relationship in the fossil record will provide a valuable benchmark and constraint on what processes can cause it.

**Highlights:**

- Studies have found that fossil and molecular diversification rates scale with time
- Such rate scaling hints at a disconnect between micro- and macroevolution
- These trends are absent from the fossil record when controlling for sampling biases
- Rate scaling in general may be artefactual, or fossils could show distinct patterns

## Results and discussion

The presence of age-rate scaling (ARS) is a long-studied phenomenon in paleobiology^1,2^, and bears close resemblance to the problem of ‘spurious’ self-correlation when a ratio is plotted against its denominator across ecology, evolution, geology and statistics^3–7^. For instance, when an evolutionary rate such as the number of extinctions per-Myr is plotted against time, a negative correlation is almost inevitable^8^. In the palaeobiological literature, rates of origination and extinction and typically measured over specified geological intervals such as geological stages. These intervals are irregular in duration, so, early on, rates were normalised for interval duration to obtain per-Myr rates. However, this is inaccurate since events are not evenly spread throughout geological stages^2,9^, might even be erroneous if diversification (origination – extinction) happens in pulses^10,11^, and propagates error in the estimation of geological intervals^1,9^. Foote showed that per-Myr rate metrics indeed suffered from negative correlation between rate and interval duration, leading to the abandonment of normalisation by interval duration in favour of per-interval rates^1,12^.

A recent study by Henao-Diaz et al. has reinvigorated interest in ARS^13^, and concluded that ARS – which might more accurately be called duration-rate scaling, as clades can occur either far back in time or near the present – was present in both diversification rates estimated from molecular phylogenetic trees and fossil time series. They compiled a dataset of first and last appearances for 90 orders of mammals, plants, and marine animals from Sepkoski’s compendium^14^. Diversification rates were calculated via Foote’s per-capita metric (See STAR Methods) and normalised by interval duration^15–17^. ARS was subsequently studied between the duration of taxonomic orders and their mean diversification rates through time. Their findings suggest that rates across different timescales cannot be compared^18^, and if true might reveal differences between micro- and macroevolutionary processes^19,20^. However, apart from the issue of plotting ratios against their denominator, a key concern with the use of Foote’s metrics is that those equations are biased by sampling and taxonomic artefacts^9,21,22^. Therefore, the question is whether ARS in fossil diversification rate analyses is caused by failure to control for sampling biases.

Here, we build on the work of Foote^1^ and Henao-Diaz et al.^13^ by investigating how different sampling artefacts might exaggerate ARS, and if they can be addressed using more appropriate methods. We do note that some palaeobiologists might inherently question the usefulness of plotting rates against clade duration, and we neither agree nor disagree with this view here. There are two steps to our analysis: (i) We gather occurrence data for marine animal orders from the Paleobiology Database (n = 408 orders after data cleaning; http://www.paleobiodb.org). Foote’s metrics are only capable of using full age-ranges (first and last appearances) of taxa, but fossil occurrence data containing age, locality and identity for each taxon is now readily available. Fossil occurrences also inherently contain information about sampling intensity through time. (ii) We compare Foote’s per-capita rates to Alroy’s second-for-third metrics (See STAR Methods)^21^ to calculate rates of origination and extinction. Second-for-third metrics require occurrences, and use a sliding-window approach that allows the examination of sampling in small time windows ^9,22^. Second-for-third metrics do not suffer from the biases that age-range metrics such as Foote’s per-capita method suffer from^21,22^. Unless stated otherwise, rates are expressed as the mean genus extinction and origination per-interval rates of an order and are not normalised for the interval duration (See STAR Methods). Rates are correlated to the duration of their respective taxonomic order, which is the time between the first and last appearance.

We find that when per-capita metrics are used, we indeed recover strong negative rate-scaling between log-transformed estimates of diversification rates and order duration (Figures 1A-1B). These relationships (origination: β = −0.52; *R*^2^ = 0.27; P < 0.001, extinction: β = −0.52; *R*^2^ = 0.27; P < 0.001) are stronger than those that had been reported previously in fossil data (e.g. origination: β = −0.23; R2 = 0.15; P < 0.001, and extinction: β = −0.25; R2 = 0.13; P < 0.01)^13^, and are of comparable strength to those reported for molecular data (e.g. speciation: β = −0.54; R2 = 0.34 and extinction: β = −0.55; R2 = 0.16)^13^. However, when we control for incomplete and heterogenous sampling by using second-for-third metrics, we find only a very weak relationship for origination (β = −0.16; *R*^2^ = 0.02; P = 0.014) and no significant relationship for extinction rate (β = −0.11; R2= 0.01; P = 0.077; Figures 1C-1D). This suggests that unequal sampling through time and between clades causes the previously-observed ARS in the fossil record.

**Figure 1.**
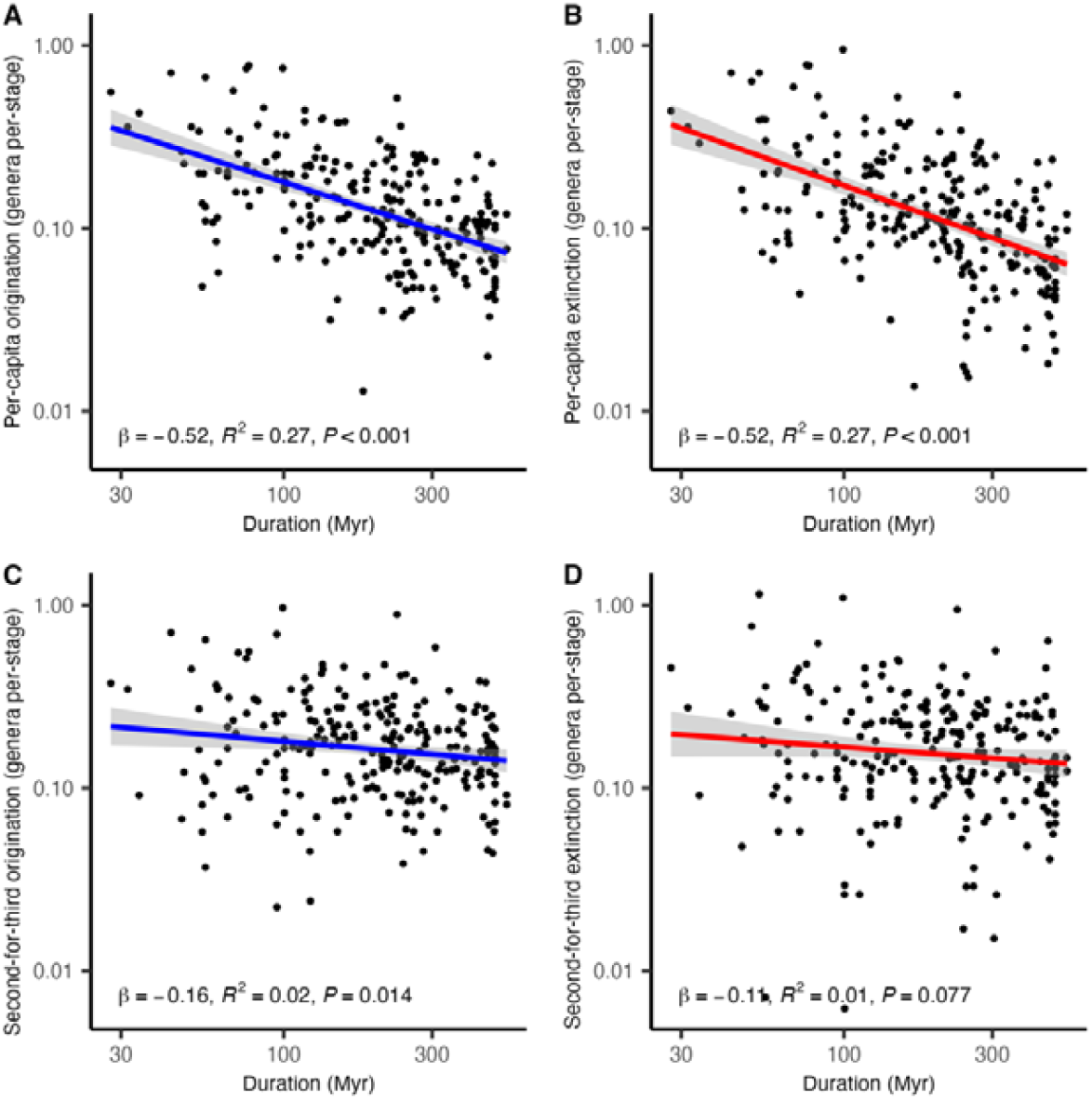
Empirical relationship between log-transformed diversification rates and clade durations. A and C show mean log-transformed genus-level origination rates (blue), and B and D show mean extinction (red) rates plotted against log clade duration for 245 marine orders sampled across the Phanerozoic. Rates in A and B are calculated via Foote’s per-capita metric, and C and D are calculated by Alroy’s second-for-third metric. Rates are shown as the mean number of events per-interval, per-lineage and not normalised for interval length. Coloured lines represent ordinary least squares regression fits between log-rate and log-duration, and shaded areas show 95% confidence intervals. β corresponds to the standardised regression coefficient that is equal to the slope of the relationship between rate and duration when their variance has been standardised to 1. Results are shown for occurrences binned in geological stages.

The occurrences that are used for the focal analysis are binned in time intervals based on geological stages. These stages vary in duration, and if a single origination and extinction pulse were to happen within a short interval, this might result in ARS. Equally, ARS in molecular phylogenetic studies tends to be strongest over the shortest time intervals (<10 Myr)^13,23^. To counteract this, we created composite time-bins with approximately equal duration of ~11 Myr by lumping the same stage-level intervals^24,25^. Even when using these equal time bins, we still find strong ARS for per-capita rates (Table S1), although both the variance explained, and the strength of ARS are decreased substantially for extinction (β = −0.39; *R*^2^ = 0.15; P < 0.001). That a strong relationship remains is not surprising, because this again shows the strong role of sampling bias. When second-for-third rates are used, ARS is again reduced. However, even under second-for-third rates, both origination (β = −0.18; *R*^2^ = 0.03; P = 0.007) and extinction (β = −0.20; *R*^2^ = 0.04; P = 0.003) now show significant trends, albeit very weak. This is consistent with the idea that when occurrences are binned in longer time intervals (effectively increasing the minimum age of orders as intervals are aggregated together), additional survivor and selection bias is applied to the clades in our dataset (see below). This highlights that heterogenous and incomplete sampling causing ARS in the fossil record arises independent of timescale and shows an additional role for selection bias.

Sampling biases are thus the likely cause of ARS in fossil time series. Drawing a parallel to phylogenetic studies, Louca et al.^23^ (also see^26^) argued that there are two distinct mechanisms that lead to a biased sample of species-rich phylogenies that can cause ARS. Mechanism 1 considers that even if clades share the same intrinsic speciation and extinction rates, clades can show a wide distribution of clade sizes because of the stochastic variation in rates through time. It is likely that clades that radiated rapidly at their inception will have survived – the “push of the past”^26,27^. This survival bias effectively truncates the distribution of clade sizes by excluding species-poor and extinct clades and can bias towards estimating higher rates, especially in young clades. Mechanism 2 suggests that diversification rates can be truly heterogenous among clades. If high rates are indeed accurately estimated, the same survival bias may, however, still apply. Additionally, selection bias may contribute to ARS under both mechanisms by only selecting clades beyond a certain species diversity-threshold, and taxonomic practices may further exaggerate this if clades are only described after reaching certain diversity. Diversity thresholds are difficult to avoid in phylogenetic studies, because extinct clades are not considered, and estimating rates for small clades can be unreliable. This again disproportionally affects younger clades, because they likely contain fewer species. Comparable selection and survivor biases could equally affect ARS in the fossil record.

One advantage of using fossil occurrences is that extinct clades are incorporated, which avoids truncating the possible diversification trajectories a clade can follow. There is also less taxonomic or researcher-focused bias in selecting clades to study that are above a certain diversity threshold, because occurrences are described per palaeontological collection and not by specifically sequencing a particular clade or species. However, this does not mean that estimating rates for small clades becomes significantly easier, and the clades that are fossilised might equally represent a biased sample of all clades that ever lived. Therefore, arbitrary diversity-thresholds are still imposed for methodological reasons, and biased preservation of the most diverse clades cannot be directly ruled out (see sections below).

We investigate how imposing diversity thresholds might influence ARS by only using orders that have accumulated ≥10 genera, in line with previous work that has recovered ARS from fossil data^13^. This analysis suggests that there is a minimal impact on observed ARS when we look at occurrences binned at the stage-level (Table S1). However, when occurrences are binned in equal-time bins, we do observe a modest but clear decrease in slope and increase in *R*^2^ (Table S1). This is likely due to the fact that very few young clades with low richness and low rates survive long enough to be sampled, because the minimum duration has increased, and few remain after filtering. Consequently, there is a minimal but observable effect of duration- and diversity-threshold selection bias on ARS.

Sampling bias in the fossil record cannot be entirely attributed to diversity-threshold sampling bias. For species to show up in the fossil record they will need to fossilise, be preserved through time, and be sampled in the present. The simplest way of thinking about sampling a species is by assigning it a probability that it will be sampled per time interval – ρ, similar to speciation and extinction rates^28,29^. Under this assumption, all else being equal, the duration of a species correlates positively with the chance that it will be sampled at least once. For higher-level taxa such as genera and orders, this also means that the more species they contain, the higher their chance of preservation. We hypothesise that especially for short duration clades there will be limited opportunities (i.e. in multiple time bins) to be sampled, which might inflate the number of false originations and extinctions.

We investigate this hypothesis with a simple heuristic simulation model of species diversification and sampling. The simulation model is based on a birth-death model with speciation and extinction^22^. Simulations are started with N species (N = 1,000 in Figure 2), and speciation and extinction occur in pulses at the boundaries between intervals (E = 0.3 in Figure 2). We further assume that the number of species is constant through time^9,30^, so that every species that is lost is immediately replaced by a new species. The sampling probability, s, of species within and between intervals is uniform. The point of this simulation is not to be exhaustive, nor to pinpoint the precise sampling artefact causing ARS, but simply to demonstrate whether incomplete sampling could conceivably cause ARS. Moreover, more complex models might induce their own artefacts leading to ARS by, for instance, inducing survivorship and selection biases^26,31^, which we seek to avoid here.

**Figure 2.**
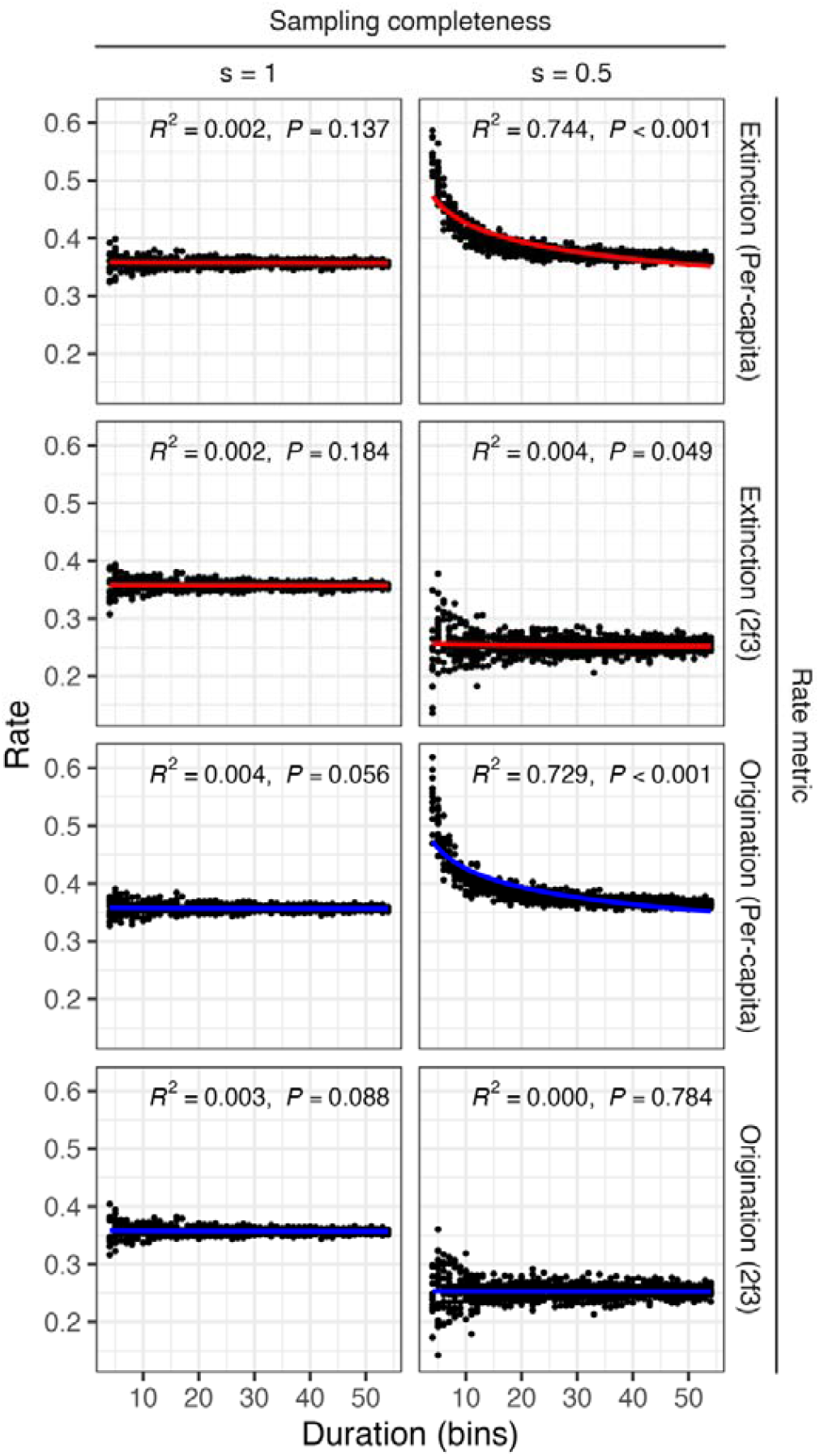
Diversification rate scaling with clade duration for simulations. Average extinction (red) and origination (blue) rates are plotted against their clade duration for 1000 repeats. Column 1 shows the relationships between rates and clade durations when sampling is complete, whereas column 2 shows a scenario of moderate sampling where each species has 50% chance of being preserved and sampled within a bin. Rows represent the rate metrics used for the empirical analysis. Row 1 and 3 use Foote’s per-capita metrics, and row 2 and 4 use Alroy’s second-for-third metrics that correct for sampling bias. Coloured lines represent non-linear least squares regression fits according to the function *y* = *ax*^*b*^. 95% confidence intervals are not shown as they overlap almost completely with the coloured lines. P-values are obtained by comparing the non-linear regression to an intercept only model. Simulations are shown for a species pool of *N* = 1000 and an extinction probability of *E* = 0.3 per interval.

We find that under these simple assumptions, ARS is absent when all species are fully sampled (s = 1), and diversification appears constant among clades irrespective of clade duration (Figure 2). However, when we impose lower sampling probabilities (s = 0.5), ARS does manifest, but only under the per-capita rates. This result is consistent under a wide range of parameter values (Figure S1). This result confirms that error in the rates for short-duration clades, caused by incomplete sampling and resulting sampling artefacts, could explain the presence of ARS in the fossil record.

Apart from data driven artefacts, statistical artefacts could cause ARS to arise in fossil data. As previously mentioned, normalising rates to per-million-year units results in ARS when rates measured over geological stages of varying durations are compared^1^, and, more generally, studies have attributed ARS to plotting a ratio (events Myr^-1^) against its denominator (Myr)^8,32^. Our results (Figure 1) demonstrate that ARS arises in per-capita rates even if they are not normalised for interval duration. However, it is possible that normalisation could either increase or weaken the strength of ARS. We find a minimal effect of normalisation on both per-capita and second-for-third rates (Table S1). These results should not be overinterpreted, because we use rates averaged over time intervals of irregular duration. The interpretation that steeper slopes equal greater independence between events and duration, or that this signals the absence of spurious self-correlation consequentially both hold little weight.

Our results emphasise the importance of controlling for heterogeneous sampling when calculating rates of origination and extinction from fossil data. This is not a new suggestion, and is true for many other metrics (e.g. species diversity^33^), but we specifically show that failing to control for incomplete sampling may induce ARS. With the increased availability of taxonomic occurrence records, and the development of sophisticated methods for estimating rates (e.g. PyRate^34^, fossilised birth-death range process^35^, second-for-third rates^21^), there is little excuse for using tools that have been shown to be erroneous^11^. Nevertheless, even when a method is chosen that can incorporate uneven sampling, care should be taken to avoid introducing artefacts relating to plotting rates against time^5^.

Various processes have been argued to cause higher rates at the start of clade radiations, with diversification slowing down with time^36^. It might therefore be surprising that controlling for uneven sampling through time significantly reduces ARS in fossil time series. Survivorship bias can cause such slowdowns and is inherently present in the fossil record^26,31^, making it an important null model for diversification patterns. Assuming constant rates of diversification, young clades will have a higher risk of extinction compared to older clades as they statistically will harbour more species. Consequentially, observable clades are likely those that diversified rapidly at their inception – the “push of the past”^26,27^. If preservation is a per-species probability^28^, clades that have experienced high origination rates are more likely to have entered the fossil record^31^. This leads to ARS, since the only short-duration clades that can be observed are those that radiated rapidly, whereas older clades regress towards their mean diversification rates. We find some support for this, because ARS increases when equal time bins and diversity thresholds are used. However, this effect is minor and when we correct for the heterogeneity, and incompleteness of sampling through time, ARS all but disappears. The absence of ARS should be further explored, and we hypothesise that a difference in taxonomic level causes this. Specifically, genera are the most common unit for fossil diversification analyses, but in contrast to species, do not go extinct until all their constituent species have gone extinct (see STAR methods).

In conclusion, we do not find strong support for a relationship between the duration of a clade and the rates of origination and extinction that it has experienced. Previous studies only found such a relationship because they used metrics that are susceptible to various sampling biases inherent to the fossil record – akin to the idea that measuring rates over shorter time scales is more error prone^37^. Our findings show that biases inherent to the fossil record may cause rate-scaling, suggesting that artefactual self-correlation^4,8,32^ is not always applicable nor that a general cause necessarily explains rate-scaling. However, the interaction between sampling, survival, and taxonomic biases provide motivation for further research to uncover the reasons underlying the different signals present in molecular^23^ and fossil data.

## Methods

### Fossil occurrence data assembly

We retrieved fossil occurrence data from the Paleobiology Database using the URL API in identical fashion to previous work^24,38^. We reiterate what has been stated there, but we performed a new download of the data on the 8^th^ of May 2024. Our analyses focus on marine Animalia. For our API calls we include all fields for marine Animalia excluding Chordata, Chordata excluding Tetrapoda, and Tetrapoda. Calls were restricted to marine environments only by the command “envtype=marine”. Separately, lists of occurrences were downloaded for Anthozoa, Brachiopoda, Bryozoa, Conodonta, Cephalopoda, Crinoidea, Decapoda, Echinoidea, Gastropoda, Graptolithina, marine Bivalvia, and Trilobita to define taxon sets/clades, and orders were defined by the data field “order”.

Diversification rate analyses were conducted at genus level, so our empirical rates pertain to the origination and extinction of genera. Occurrences were filtered to only those that were assigned with certainty to at least genus-level using “idqual=genus_certain” in our calls. We only took into account the latest identification (“idtype=latest”) of the occurrence, only valid taxa (“taxon_status=valid”), and removed form taxa (e.g., assigned based on morphological affinity but not phylogeny) or ichnotaxa (e.g., trace fossils) via “pres=regular”. We further filtered the occurrences to those that had been assigned to a specific order.

Occurrence were binned via two schemes: (i) 100 geological stages, and (ii) 49 composite equal time bins. The equal time bins each spanning approximately ~11 myr were created by binning geological stages^39^. We assigned the occurrences to bins based on a majority rule. If a time bin contained >50% of the geological time range associated with the occurrence, it was assigned to that bin^39^. Our downstream diversification rate calculations depend on clades ranging over at least 3 time bins. Any order with fewer bins was removed from the analysis. After applying these sifting criteria, the two final datasets based on geological and composite time bins consisted of respectively 610,514 and 646,204 occurrences, 408 and 368 orders, a clade duration range of 5.45-520.99 and 31.60-538.79 Myr and face-value diversity counts ranging from 1 to 1267 and 1 to 1299 genera per order.

### Estimating diversification rates

The quantification of changing rates of origination and extinction has been one of the central goals of palaeobiology. A substantial number of metrics have been developed over time, and have been scrutinised equally as often^1,22,33^. Alroy^15^ and Foote^16,17^ laid the foundation to modern rate metrics that quantify how rates change between subsequent time intervals, i.e. geological stages and equal time bins, whereas later iterations and metrics have built on this framework^9,22,30,40^. Here, we explore the effect of Foote’s^17^ per-capita rate metric and Alroy’s^40^ second-for-third metric on the estimation of diversification rates and the impact on rate-age-relationships.

Foote’s metrics rely on the age ranges of organisms, i.e. their first and last appearance in the fossil record. Four types of ranges are classified: taxa present both before and after a focal interval, crossing its base and top (Nbt); taxa present before and within the interval but not after, crossing only its base (NbL); taxa present in and after the interval, crossing only its top; and taxa found only within the interval (NFL). Summed, these counts are the range-through diversity (Nbt). The rates we use here are Foote’s per-capita rates where origination is 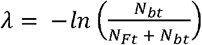 and extinction is 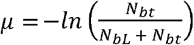 Note that Foote’s metrics were originally normalised by the interval duration over which they were measured^17^, but this was later reconsidered as discussed below^12^. These range based metrics are still actively used^41^, and are perhaps the best approach to study range based data^33^. However, range-based metrics have been argued to suffer from edge, Pull of the Recent, and Signor-Lipps effects, and taxonomic biases. In particular, this means that extinction rates are increased in intervals preceding mass extinctions (effectively smeared out), rates gradually decline as the recent is approached, and the importance of rare taxa is overweighted when sampling is good (i.e. Pull of the Recent) or underweighted when extinction is fast (i.e. Signor-Lipps)^22,40^.

Fossil occurrence data has become increasingly available, promoting a way to move beyond using range-based metrics and focussing on patterns of taxonomic sampling throughout. Alroy’s second-for-third metric has been argued to not suffer from any of the above biases, and relies on a sliding-window approach, in contrast to full ranges^40^. Our focal time interval is *i*_0_, and we assume a cohort of taxa that have only been sampled since *i*-_1_ up to *i*_2_ for extinction and *i*-_2_ to *i*_1_ for origination. The following counts are of importance for extinction (origination in brackets if distinct): *s*_1_ = taxa sampled in *i*-_1_ and *i*_0_ (*i*_0_ and *i*_1_) but no later (earlier), *s*_2_ = taxa sampled in *i*-_1_ and *i*_1_ (*i*-_1_ and *i*_1_) but not in *i*_0_ or *i*_2_ (*i*-_2_ and *i*_0_), *s*_3_ = those sampled in *i*-_1_ and *i*_2_ (*i*-_2_ and *i*_1_) but not in *i*_0_ or *i*_1_ (*i*-_1_ and *i*_0_), *t*_2_ = those sampled in *i*-_1_ and *i*_0_ (*i*_0_ and *i*_1_) regardless of being sampled later (earlier), and p = *i*-_1_ and *i*_1_ but not in *i*_0_. The formula for extinction and origination proportion (not rate) is then calculated as *O* = *E* = (*s*_1_ − *s*_*x*_)/(*t*_2_ + *p*). If sampling is uniform and given that unextinction cannot occur the counts should have the expected order *s*_1_≥ *s*_2_ ≥ *s*_3_, and *s*_*x*_ would be *s*_3_ (the Gap-Filler equation^22^). However,in practice they do not and might even show the reverse pattern *s*_1_ < *s*_2_ < *s*_3_. To address this Alroy suggested algorithmically replacing *s*_*x*_ with the second lowest of the counts, e.g. *s*_*x*_ = *s*_1_ if *s*_2_ > *s*_1_ > *s*_3_. Conventional rates (e.g. decay coefficients) are then obtained by log-transformation where 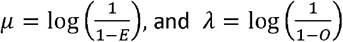 Second-for-third rates are obviously heuristic because of the, but it incorporates as much information as possible to correct for sample size.

We focus on genus origination and extinction per-interval rates for both Foote’s and Alroy’s metrics, and do not normalise for interval duration. That is, we focus on the rate at which species originate and go extinct per taxon per interval, and do not normalise our rates to be expressed per million years as was originally done^17^. Raup and Sepkoski, although using percentage extinction and not true rates, already discussed preferring per-interval metrics in their seminal paper^2^. They observed that the number of raw extinction counts did not correlate with interval duration as they are likely clustered in time^9,33^. Foote further explored the consequences of normalisation and found that if intervals were to experience a single peak in extinction risk, this would dominate the total extinction profile of short intervals more than for long intervals, effectively causing a negative correlation between interval duration and extinction rate^1^. We repeat our analysis for time-normalised rates to understand if negative rate scaling between bins has an effect on average rate comparisons between clades.

We calculate both Foote’s per capita rates and second-for-third rates using the R package *divDyn*^42^. Origination and extinction rates were calculated for all orders for both geological stages and equal time bins, and then averaged per order. In some instances, an average rate of zero is estimated. Rate estimates of zero likely are caused by sampling error or model misspecification and make little sense empirically^33^ (but see ref^43^). For example, an order must have had a positive origination rate if it contains more than one species. In our focal analysis, orders with average rate estimates of zero are removed (i.e. the number of orders is reduced from 408 to 245 for stage-level bins and from 368 to 220 for equal-time bins). This might bias the analysis towards observing greater rate-scaling. Regardless, average rates are subsequently correlated across all orders with their duration using (i) non-linear least squares, and (ii) linear regression by log-transforming both rate and duration, which would have removed zero estimates.

### A simulation model

We investigate if incomplete sampling through time can cause ARS with a simulation model of species diversification and sampling. The simulation model is based on a prior model used in the testing of rate metrics^22^, and can be abstracted to modelling temporal change in species identities (i.e. faunal turnover) in a species pool. The species pool has constant species diversity, *N*, through time (*N* = 10^2^, 10^3^, 10^4^), which can be justified by the strong correlation between per-interval extinction rate followed up by high origination in the next interval, across the Phanerozoic marine fossil record^9,30^. Because we are not interested in measuring diversity, a variable trend might have sufficed as well. At the end of each interval, species have a probability, *E*, to go extinct (*E* = 0.15, 0.3, 0.6). Speciation or origination is modelled as the replacement of the extinct species immediately upon extinction by a new species. Effectively, diversification thus occurs in pulses between intervals. We simulate 10^3^ clades per parameter combination to calculate ARS, with clades experiencing faunal turnover for a random number of intervals drawn from a uniform distribution ranging from 3 (i.e. the minimum number of bins required to calculate per-capita rates) to 54 (i.e. roughly the reflecting the length of the Phanerozoic if each interval is 10 Myr) intervals. The realised time series of faunal turnover, each seen as a unique clade, are then degraded based on a per-species sampling probability per interval, *s*. We model sampling intensity from poor to good (*s* = 0.25, 0.5, 0.75). The complete and degraded time series are then used to calculate both Foote’s per-capita and Alroy’s second-for-third rates and plotted against the number of time intervals to repeat the empirical analysis (Figures 3, S5, and S6).

Our simulations assume that sampling is uniform through time and across clades. In practice, sampling within time bins will vary because of variation in preservation (e.g. substrate), sampling effort, and fossilisation potential of clades (e.g. soft or hard-bodied). Taxonomic biases might further contribute to error in the estimation of rates by over or under splitting taxa. This might in itself result from incomplete sampling through time, because infrequently sampled but temporally long-ranged taxa might be under split^22^. Equally, taxonomic ranks are inherently arbitrary concepts. Especially when it comes to orders and other higher ranks, it might be difficult to accurately delineate when they arise, and consequently higher diversity orders might be recognised before low diversity orders. Therefore, our simulations are by no means exhaustive in terms of the possible ways that sampling could influence the estimation of diversification rates.

### Genus versus species-level diversification

In the main text we discuss how survival bias might cause a “push of the past” under which clades that radiate rapidly early in their duration are more likely to be observed in the fossil record. This could lead to a strong negative relationship between diversification rate and clade duration even when controlled for heterogenous and incomplete sampling, but we do not find this. An explanation for this absence could be our empirical focus on fossil genera instead of species. This difference in taxonomic level, often required in fossil analyses, becomes apparent when the survival of a species (i.e., species duration), is compared to that of a genus. If a species’ extinction rate is constant, the probability of species survival decreases linearly through time^44^. In contrast, if we look at genera, which are each made up of one or more species, a genus will only go extinct after its last species disappears. With time, as a genus accumulates more species, the survivorship of the genus therefore increases^45–47^. This could be modelled by assuming a small probability of genus formation at each speciation event, and that each genus starts as a single species from which it diversifies. Consequentially, the rate of genus origination would correlate positively with the rate of species formation, albeit stochastic. If a “push of the past” effect occurred on the species-level, a likely identical preservation bias for species and genera would also operate. However, because our results do not reveal ARS, the formation of genera could be largely independent of the formation and extinction of species. We speculate that this might consequently be caused by taxonomic or biological reasons.

From a taxonomic perspective, evolutionary lineages in the fossil record might not be recognised as independent genera until they have shown sufficient morphological differentiation. This might cause different rate-scaling patterns at the level of species (molecular data) and genera (fossil data) to arise, especially if we consider the prevalence of cryptic species in molecular datasets that show little morphological divergence but are reproductively isolated^48^. If most of these cryptic species are unlikely to persist over evolutionary time, which could be the case in some contemporary clades such as sticklebacks^49^, we might expect the highest rates in young clades with a substantial proportion of cryptic species. If genera or species are delineated by morphology in the fossil record, such cryptic diversity would only be detectable in molecular datasets.

From a biological perspective, the assumption that a genus has a survival probability, and thus preservation probability, that is a function of the number of species contained could be relaxed. For instance, even if a genus is species-rich, these species might all be extinction-prone because of a property such a specialisation or range size, whereas a genus with a single species might consist of a generalist that is widespread. There are thus two ways in which clades with few genera might have a high survival probability through time. Either each of these genera are widespread, or if the number of species matters, few genera might contain many species purely by chance or taxonomic practice. A push of the past at the level of species may consequentially not be experienced at the level of the genus, either because of stochasticity, the extinction vulnerability of the species within the genus, or by taxonomic relationships not being observable in the fossil record. However, the interaction between these processes is complex and future studies might benefit from extended null models to analyse these patterns.

### Quantification and statistical analysis

All statistical analyses are described above (Method details). Statistical analyses and simulations were conducted in R 4.3.1^50^.

## Acknowledgements

We thank Alex Pigot and David Murrell for discussion and encouragement. Three anonymous reviewers greatly improved the quality of the paper. The Paleobiology Database enabled this work, and contributors are wholeheartedly thanked for their effort. B.R.R. and R.A.C. are funded through the Royal Society by URF\R1\211571 awarded to R.A.C. This is Paleobiology Database publication XXX.

## Author contributions

B.R.R. conceptualised the study. R.A.C. curated the data. B.R.R. designed, programmed, and conducted the analysis. B.R.R. wrote the initial draft, which all authors edited and approved.

## Declaration of interests

The authors declare no competing interests

## Notes

### Competing Interest Statement

The authors have declared no competing interest.

### Summary of Updates

Updated main text and supplement. No change in results nor conclusions.

