## Supplemental information Figure S1 and Table S1 for "Apparent timescaling of fossil diversification rates is caused by sampling bias"

**Table S1. Ordinary Least Squares regression results for the five scenarios explored.**
OLS regressions were fit between log-transformed mean rates and durations of orders. β corresponds to the standardised regression coefficient that is equal to the slope of the relationship between rate and duration when their variance has been standardised to 1. PC and 2f3 respectively refer to the Foote's per-capita and Alroy's second-for-third rate metrics.

| Rate | Metric | β | β-S.E. | t-value | P-value | $R^2$ |
|---|---|---|---|---|---|---|
| **1) Per-interval rates – geological stages (all clades) N = 245** | | | | | | |
| Origination | PC | -0.517 | 0.055 | -9.422 | < 0.001 | 0.268 |
| Extinction | PC | -0.518 | 0.055 | -9.448 | < 0.001 | 0.269 |
| Origination | 2f3 | -0.157 | 0.063 | -2.475 | 0.014 | 0.025 |
| Extinction | 2f3 | -0.113 | 0.064 | -1.777 | 0.077 | 0.013 |
| **2) Per-interval rates – equal time bins (all clades) N = 220** | | | | | | |
| Origination | PC | -0.544 | 0.057 | -9.58 | < 0.001 | 0.296 |
| Extinction | PC | -0.391 | 0.062 | -6.27 | < 0.001 | 0.153 |
| Origination | 2f3 | -0.181 | 0.067 | -2.719 | 0.007 | 0.033 |
| Extinction | 2f3 | -0.198 | 0.066 | -2.989 | 0.003 | 0.039 |
| **3) Per-interval rates – geological stages (clade richness ≥ 10) N = 226** | | | | | | |
| Origination | PC | -0.538 | 0.056 | -9.54 | < 0.001 | 0.289 |
| Extinction | PC | -0.53 | 0.057 | -9.353 | < 0.001 | 0.281 |
| Origination | 2f3 | -0.149 | 0.066 | -2.258 | 0.025 | 0.022 |
| Extinction | 2f3 | -0.135 | 0.066 | -2.047 | 0.042 | 0.018 |
| **4) Per-interval rates – equal time bins (clade richness ≥ 10) N = 207** | | | | | | |
| Origination | PC | -0.619 | 0.055 | -11.287 | < 0.001 | 0.383 |
| Extinction | PC | -0.431 | 0.063 | -6.835 | < 0.001 | 0.186 |
| Origination | 2f3 | -0.207 | 0.068 | -3.029 | 0.003 | 0.043 |
| Extinction | 2f3 | -0.238 | 0.068 | -3.51 | < 0.001 | 0.057 |
| **5) Normalised rates – geological stages (all clades) N = 241** | | | | | | |
| Origination | PC | -0.463 | 0.057 | -8.074 | < 0.001 | 0.214 |
| Extinction | PC | -0.543 | 0.054 | -9.998 | < 0.001 | 0.295 |
| Origination | 2f3 | -0.138 | 0.064 | -2.159 | 0.032 | 0.019 |
| Extinction | 2f3 | -0.207 | 0.063 | -3.275 | 0.001 | 0.043 |

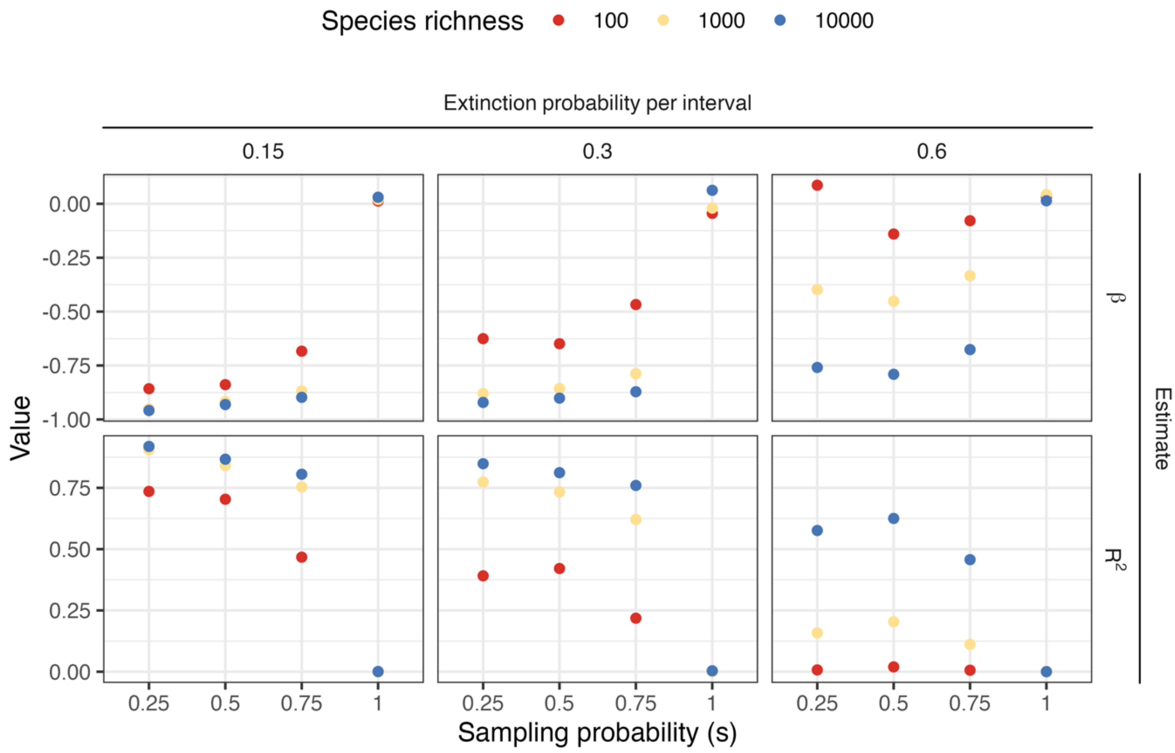


**Figure S1. Parameter and $R^2$ estimates for age-rate scaling relationships in the simulations.** Clades were simulated under a simple birth-death model in which pulses of extinction and origination occur between time intervals. Results shown are for OLS fits to log-log relationships between per-capita extinction rate and clade duration. Columns represent varying probabilities of extinction per interval, whereas rows show the beta coefficient and $R^2$ values. On the x-axis of each facet the sampling probabilities (s) are shown, which indicates the probability that an individual species is sampled within each time interval. Relationships are calculated over 1000 simulated clades.
